# Genetic context drives age-related disparities in synaptic maintenance and structure across cortical and hippocampal neuronal circuits

**DOI:** 10.1101/2023.07.27.550869

**Authors:** Sarah E. Heuer, Emily W. Nickerson, Gareth R. Howell, Erik B. Bloss

## Abstract

The disconnection of neuronal circuits through synaptic loss is presumed to be a major driver of age-related cognitive decline. Age-related cognitive decline is heterogeneous, yet whether genetic mechanisms differentiate successful from unsuccessful cognitive decline through synaptic structural mechanisms remains unknown. Previous work using rodent and primate models leveraged various techniques to suggest that age-related synaptic loss is widespread on pyramidal cells in prefrontal cortex (PFC) circuits but absent on those in area CA1 of the hippocampus. Here, we examined the effect of aging on synapses on projection neurons forming a hippocampal-cortico-thalamic circuit important for spatial working memory tasks from two genetically distinct mouse strains that exhibit susceptibility (C57BL/6J) or resistance (PWK/PhJ) to cognitive decline during aging. Across both strains, synapses on the CA1-to-PFC projection neurons appeared completely intact with age. In contrast, we found synapse loss on PFC-to-nucleus reuniens (RE) projection neurons from aged C57BL/6J but not PWK/PhJ mice. Moreover, synapses from aged PWK/PhJ mice but not from C57BL/6J exhibited morphological changes that suggest increased synaptic efficiency to depolarize the parent dendrite. Our findings suggest resistance to age-related cognitive decline results in part by age-related synaptic adaptations, and identification of these mechanisms in PWK/PhJ mice could uncover new therapeutic targets for promoting successful cognitive aging and extending human health span.

## INTRODUCTION

Synapses are the fundamental computational subunits of the brain, and the building blocks of neuronal circuits that vary in complexity based on brain region and mammalian organism (1, 2). Synapses enable cell-type specific forms of communication and are often targeted to defined spatial subcellular locations on postsynaptic cell types (3), which constrains neural circuit operations (4, 5). Synapses act as plastic, tunable biochemical and electrical signaling compartments, a feature that allows internal states to shape neural circuit function (6, 7), and may be the primary substrate in the nervous system mediating learning and memory recall (8, 9).

Individual excitatory synapses on many neuronal cell types take the form of small protrusions along the dendrites (i.e., dendritic spines (10)), which can be visualized by multiple imaging methods (8, 11, 12). Spine structure reflects the functional strength of each connection: spine head size correlates with synapse strength (8, 12), and the morphology of the spine neck determines the degree of electrical and biochemical filtering that occurs between the synapse and the parent dendritic cable (13, 14). During normal aging, mammalian organisms experience progressive, yet variable, cognitive decline, even in the absence of a disease, which corresponds to a loss in synaptic plasticity(15). Relative to other synapses, spine synapses seem to be the primary type that are lost or altered during aging, and these changes in the aging brain appear circuit specific (16, 17). For example, the density of spine synapses decrease over time on cortical neurons in the frontal but not visual cortex, and decrease on CA3 pyramidal neurons but not on neighboring neurons in CA1 (16, 18, 19). The loss of synapse number and the changes in synaptic structure on frontal cortical neurons correlate with progressive decreases in cognitive function with age (20–22), yet reports of the degree to which synaptic loss occurs suggests significant heterogeneity (16, 23–25). The causal factors that underlie this variability could be what determines successful versus unsuccessful neural aging (16, 26, 27), and there is significant interest in understanding how genetic components and environmental influences coordinately shape the trajectory of age-related cognitive changes and synaptic connectivity.

Non-human primates (NHPs) (22, 28, 29) and rodents (rats (25, 30, 31) and mice (32)) are commonly used model systems to study the effect of aging on synapses. Although NHPs are the most translatable model for relating age-related neural circuit changes to cognitive decline (22, 33), they have long lifespans (>30 years), require complex housing systems, and are not amendable to genetic-based experimentation at scale. However, to a large extent, data from rodent models recapitulate the magnitude of age-related synaptic loss in vulnerable brain regions (including the prefrontal cortex) that has been observed in NHPs (24, 25, 34). Studies in mice have shown loss in synaptic plasticity and number during aging and in age-related diseases (35, 36). Unlike NHP experiments, which necessarily use genetically diverse non-inbred subjects, the work in mice has largely been performed using the inbred C57BL/6J (B6) mouse strain. The lack of genetic diversity limits translatability of mouse work to humans, and led us to evaluate the potential of incorporating genetically distinct wild-derived inbred mouse strains in our own work (37). Our previous results show wild-derived strains exhibit diverse cognitive and behavioral phenotypes during aging and when carrying Alzheimer’s disease (AD)-linked transgenes, suggesting genetic background strongly influences neuronal circuitry and function (38, 39). In particular, the PWK/PhJ (PWK) mouse strain did not show amyloid-dependent cognitive decline or synaptic loss, suggesting PWK is an ideal strain to study resilience to AD (40). However, the neuronal circuits vulnerable in aging are different than those vulnerable to AD, and it remains unknown whether PWK is also resistant to normal age-related synaptic loss.

To determine the effect of genetic context on age-related synapse dynamics (and how they might relate to their observed cognitive susceptibility/resilience (38, 41)), we examined aging-related synapse loss in the traditionally studied B6 mouse strain and the wild-derived PWK. PWK mice differ from B6 by approximately 20 million single nucleotide polymorphisms (SNPs) and structural variants (37). To target our analyses to specific projection neurons, we leveraged retrograde viral strategies to label two circuits known to have different vulnerabilities to aging or AD: PFC-projecting neurons to nucleus reuniens (RE; PFC-to-RE), and CA1-projecting neurons to PFC (CA1-to-PFC) (17). Spine synapses on CA1-to-PFC neurons showed marked stability throughout aging in both B6 and PWK mice. In contrast, PFC-to-RE neurons were susceptible to age-related spine loss in B6 but not PWK. Further, there were morphological changes suggestive of synapse strengthening in PWK but not B6 mice. Collectively, these results provide evidence that PWK are a model of successful synaptic aging.

## RESULTS

### Labeling strategies to visualize synapses from two neural circuits in genetically distinct mice

Our previous studies suggest PWK are a model of cognitive, neuronal and synaptic resilience to Aβ pathology (38, 40). However, since age is the leading risk factor for developing AD, we tested the prediction that PWK would also be a model of successful synaptic aging when compared to traditionally used inbred B6 mice. We leveraged previously established viral approaches (42, 43) using two different fluorophores to label excitatory projection neurons forming two arms of a connected hippocampal-cortico-thalamic loop important for spatial working memory tasks (44). Each set of projection neurons expressed either EGFP or tdTomato across cohorts of female B6 and PWK mice ranging from 4-29 months of age (see **Table S1**). Three unilateral intracranial injection sites were targeted to deliver recombinant adenoassociated virus (AAV, see **Methods**) to drive EGFP in PFC neurons that project to RE (PFC-to-RE) and tdTomato expression in CA1 neurons that project to PFC (CA1-to-PFC). After allowing 3-4 weeks for proper retrograde movement and fluorophore expression each mouse was perfused, coronal slices were made with a vibratome, slices containing EGFP+ or tdTomato+ neurons were imaged, and dendritic spine density and morphologies were analyzed (**Figure 1A**). Three age groups were assessed: young (4-7 months of age), middle-aged (11-17 months of age) and aged (22-30 months of age) groups. Neurons making up the two projection circuits in B6 and PWK mice were labeled with equivalent efficiency, suggesting a similar overall density of neurons comprising these pathways from the compared strains (**Figure 1B-C**).

**Figure 1:**
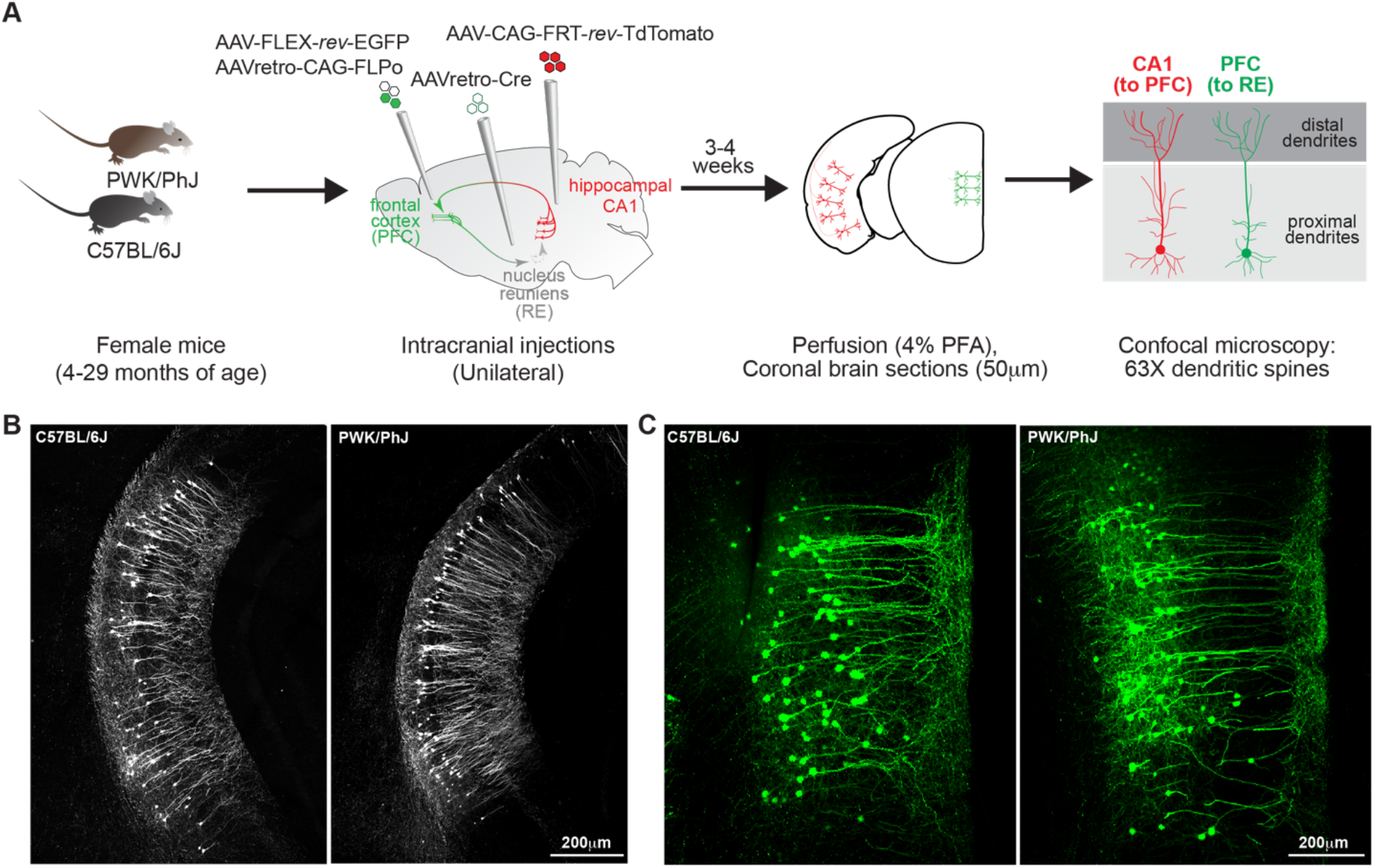
Viral labeling of PFC-to-RE and CA1-to-PFC neurons across C57BL/6J and PWK/PhJ mice. **(A)** Experimental outline (see ***Methods*** for additional details) **(B-C)** Example 10X images of tdTomato+ CA1 neurons (B) from B6 (left) and PWK (right) demonstrating consistent CA1-to-PFC projection circuit labeling, and EGFP+ PFC neurons (C) from B6 (left) and PWK (right) demonstrating consistent PFC-to-RE projection circuit labeling. Summary metadata for individual mice used in this study reported in **Table S1**.

### Aging spines on proximal CA1 dendrites exhibit stable densities but altered morphologies in PWK but not B6 mice

The CA1 region of the hippocampus is well-studied on account of its anatomy, physiology, role in episodic memory, and vulnerability to both age-and AD-related pathological insults (27). While generally thought to be relatively resistant to normal age-related synaptic loss (45–47), it is unclear if age-related structural changes occur on specific CA1 projections, and if they are modulated in subjects from genetic contexts that exhibit resistance to cognitive aging. The majority of excitatory inputs onto CA1 pyramidal cells are made onto the proximal apical oblique and basal branches (48, 49) which both receive inputs from hippocampal area CA3 (50). Therefore, we first analyzed dendritic spine synapses on branches from this proximal compartment specifically on CA1-to-PFC neurons (**Figure 2A**). Consistent with the prediction from previous reports (17, 45), we observed no significant age-related changes in proximal spine density in either B6 or PWK mice (**Figure 2B-C**).

**Figure 2:**
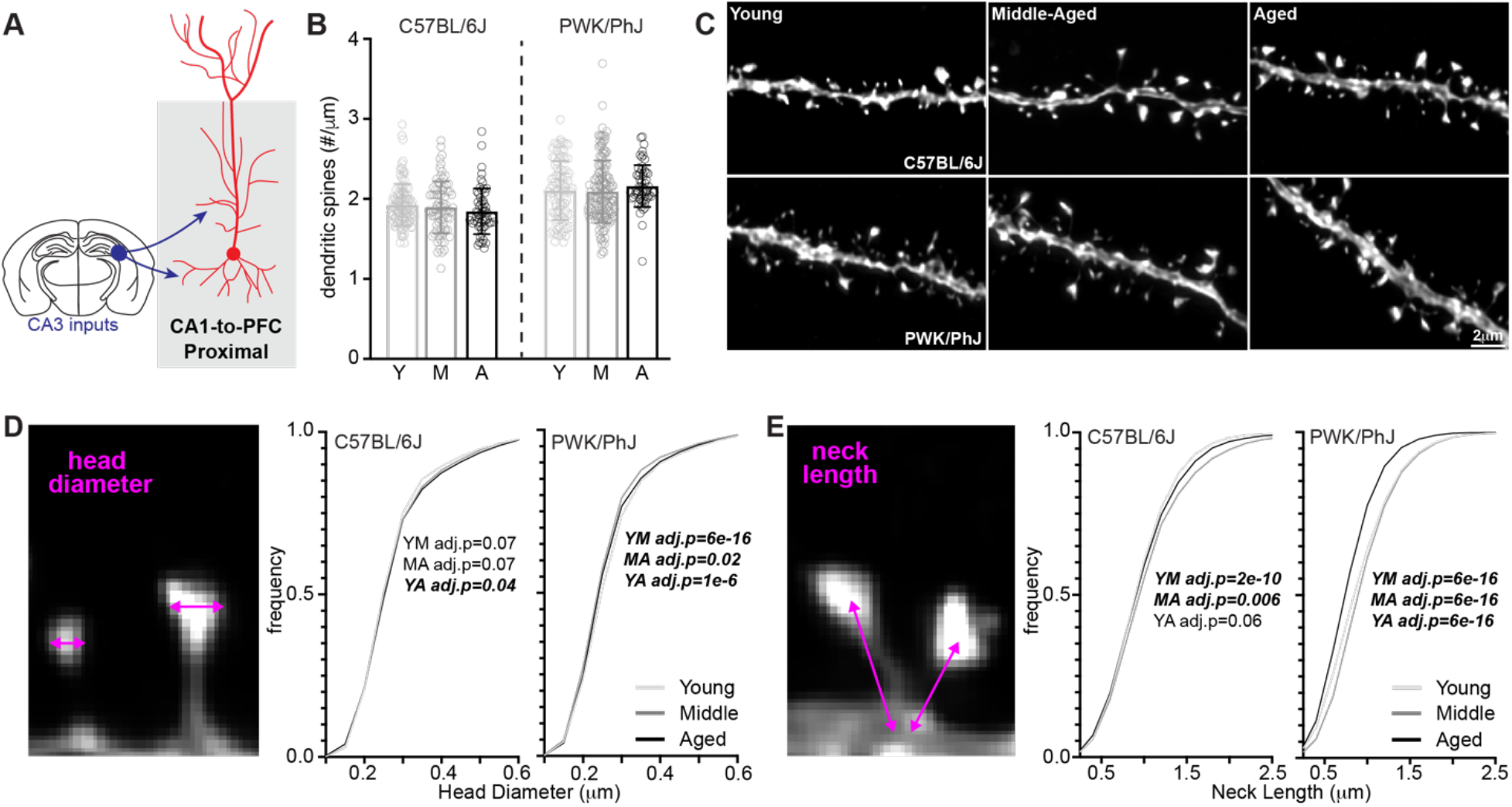
CA1-to-PFC proximal dendrites are resistant to synaptic density changes across both strains, but PWK/PhJ spines change morphologies. **(A)** Example schematic of tdTomato+ proximal CA1-to-PFC dendrites analyzed here and projection input origins in hippocampal area CA3. **(B)** Proximal CA1-to-PFC dendrite spine densities (spines/μm) comparing young (Y), middle (M) and aged (A) within each strain. Data points represent individual branches (n=20/mouse); error bars are ± SD; one-way ANOVA identified no significant effects within B6 or PWK (**Table S2**). **(C)** Representative 63X images of proximal CA1-to-PFC dendrites from each strain/age group. **(D)** CA1-to-PFC proximal spine head diameter example image (left) and cumulative distributions from B6 (middle) and PWK (right), across age groups. Kolmogorov-Smirnov tests were used to evaluate statistical significance (Bonferroni adj. p<0.05) of pairwise comparisons: young vs middle (YM), middle vs aged (MA), and young vs. aged (YA), and p-values reported on each graph (see **Table S2**). Data points are representative of measures from individual spines. **(E)** Same as **(D)** for CA1-to-PFC proximal spine neck length. Summary statistics for data points represented in each graph reported in **Table S2.**

Dendritic spines alter their morphologies in response to repeated artificial synaptic stimulation (6, 9), under conditions of homeostasis (7, 51), and in reaction to disease pathologies and during aging (52, 53). Spine head morphology is a reliable predictor of synaptic stability and strength (8, 12), so we analyzed the maximum head diameter of each reconstructed spine. Our analysis found no overt age-related changes in B6 mice, but the same analysis in PWK mice revealed a subtle yet significant age-related shift to spines with smaller head diameters (**Figure 2D**). This effect was evident when we examined the densities of spines across the smallest (Q1) and largest (Q4) quartiles of head diameters (**Figure S1B**).

The morphology of dendritic spine necks (e.g. neck width and length) are similarly plastic (13) and define the compartmentalization of spines much more critically than spine head size (13, 14, 54, 55). Dendritic spine necks act as strong electrical resistors, filtering currents from the synapse to the parent dendrite branch (54). Necks also contain receptors for neuromodulatory systems that regulate local excitability (56). Spine necks are often below 50 nm (49) making them hard to resolve with light microscopy techniques, thus we analyzed putative spine neck lengths to determine whether this aspect of spine neck morphology is readily modified during aging. Neck lengths in B6 mice were slightly lengthened with age, complementing the stability observed in spine heads. In contrast, PWK spine neck lengths showed a biphasic response, first increasing from young to middle-aged, then decreasing significantly in aged mice compared to the younger groups (**Figure 2E**, **Figure S1C**).

### Spine changes on distal CA1 dendrites mirror those found on proximal compartments

Distal tuft dendrites in CA1 pyramidal cells are thought to be the most vulnerable to AD-related synaptic changes due to their afferent inputs originating in the EC (**Figure 3A**) (57), where associated neuropathology is observed very early in the disease process (58, 59). Synapses on distal dendrites are understudied relative to those on more proximal compartments due to limitations in traditional techniques used to evaluate plasticity at remote locations relative to the soma. Consistent with our results from the proximal dendrites, we observed no significant age-related changes to distal tuft spine densities in B6 or PWK mice (**Figure 3B-C**). Like spines from proximal branches, we observed few age-related changes in head diameter and neck length in spines from B6 distal branches (**Figure 3D-E, Figure S2B-C**). However, spine head diameters were larger in aged PWK mice compared to young and middle-aged cohorts (**Figure 3D**, **Figure S2B**), and spine neck lengths were significantly shorter in aged compared to young and middle-aged mice (**Figure 3E**, **Figure S2C**).

**Figure 3:**
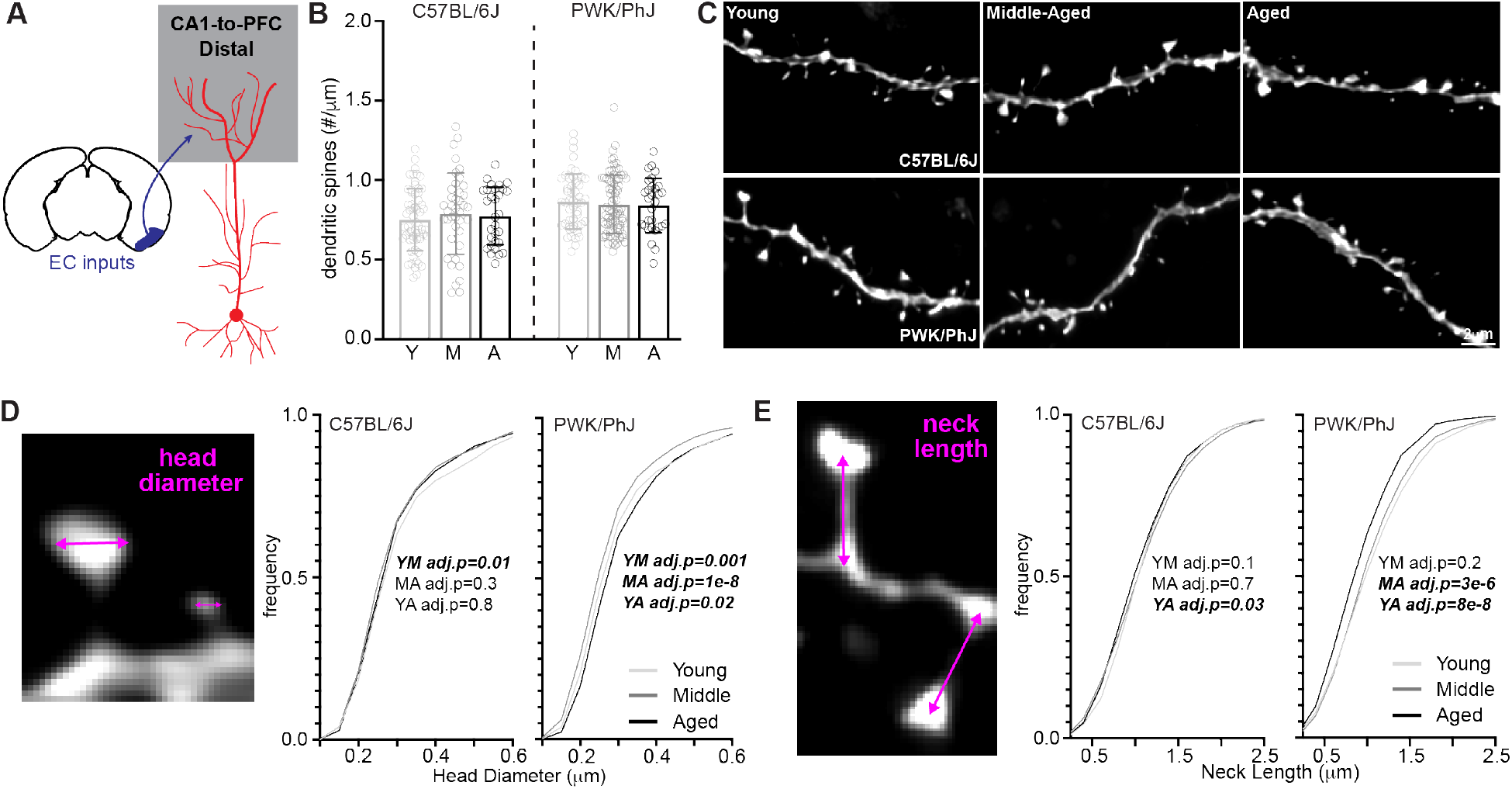
CA1-to-PFC distal tuft dendrites do not lose synapses but are morphologically altered in PWK/PhJ mice. **(A)** Example schematic of tdTomato+ distal tuft CA1-to-PFC dendrites analyzed here and projection input origins in entorhinal cortex (EC). **(B)** Distal tuft CA1-to-PFC dendrite spine densities (spines/μm) comparing young (Y), middle (M) and aged (A) within each strain. Data points represent individual branches (n=10/mouse); error bars are ± SD; one-way ANOVA identified no significant effects within B6 or PWK (**Table S3**). **(C)** Representative 63X images of distal tuft CA1-to-PFC dendrites from each strain/age group. **(D)** CA1-to-PFC distal tuft spine head diameter example image (left) and cumulative distributions from B6 (middle) and PWK (right), across age groups. Kolmogorov-Smirnov tests were used to evaluate statistical significance (Bonferroni adj. p<0.05) of pairwise comparisons: young vs middle (YM), middle vs aged (MA), and young vs. aged (YA), and p-values reported on each graph (see **Table S3**). Data points are representative of measures from individual spines. **(E)** Same as **(D)** for CA1-to-PFC distal tuft spine neck length. Summary statistics for data points represented in each graph reported in **Table S3.**

Collectively, these results show spine densities on proximal and distal dendrites from CA1- to-PFC pyramidal cells do not change with age in both B6 and PWK mice. Furthermore, B6 spines remained morphologically stable whereas PWK spines exhibited changes indicative of increased excitability at the synapse and to the parent dendritic cable, which support the notion that these adaptations could play a role in maintaining information processing in the aging hippocampus.

### Differential vulnerability to age-related synaptic changes on PFC-to-RE neurons across B6 and PWK mice

Previous studies in humans, NHPs, rats and mice report that pyramidal neurons in the prefrontal cortex are susceptible to synaptic loss with age (21, 22, 25, 29, 60). Moreover, preventing this degeneration could be key to promoting successful cognitive aging (21, 61, 62). To test the prediction that PWK mice are resistant to age-dependent synaptic loss in the PFC, we used AAVs to also label deep-layer neurons (in the exact mice described above) in the PFC that project to the RE (**Figure 1**) that appear to mediate behavioral flexibility and working memory (63–65). Like the CA1-to-PFC neurons, PFC-to-RE neurons have a similar proximal-distal dendritic architecture, and each compartment receives a distinct set of inputs. Basal and apical oblique dendrites located in the proximal regions receive afferent inputs from CA1 (among other regions), whereas distal tuft receive a distinct set of inputs from the thalamus (43) (**Figure 4A**). Therefore, we analyzed the spines on proximal and distal dendrites separately.

**Figure 4:**
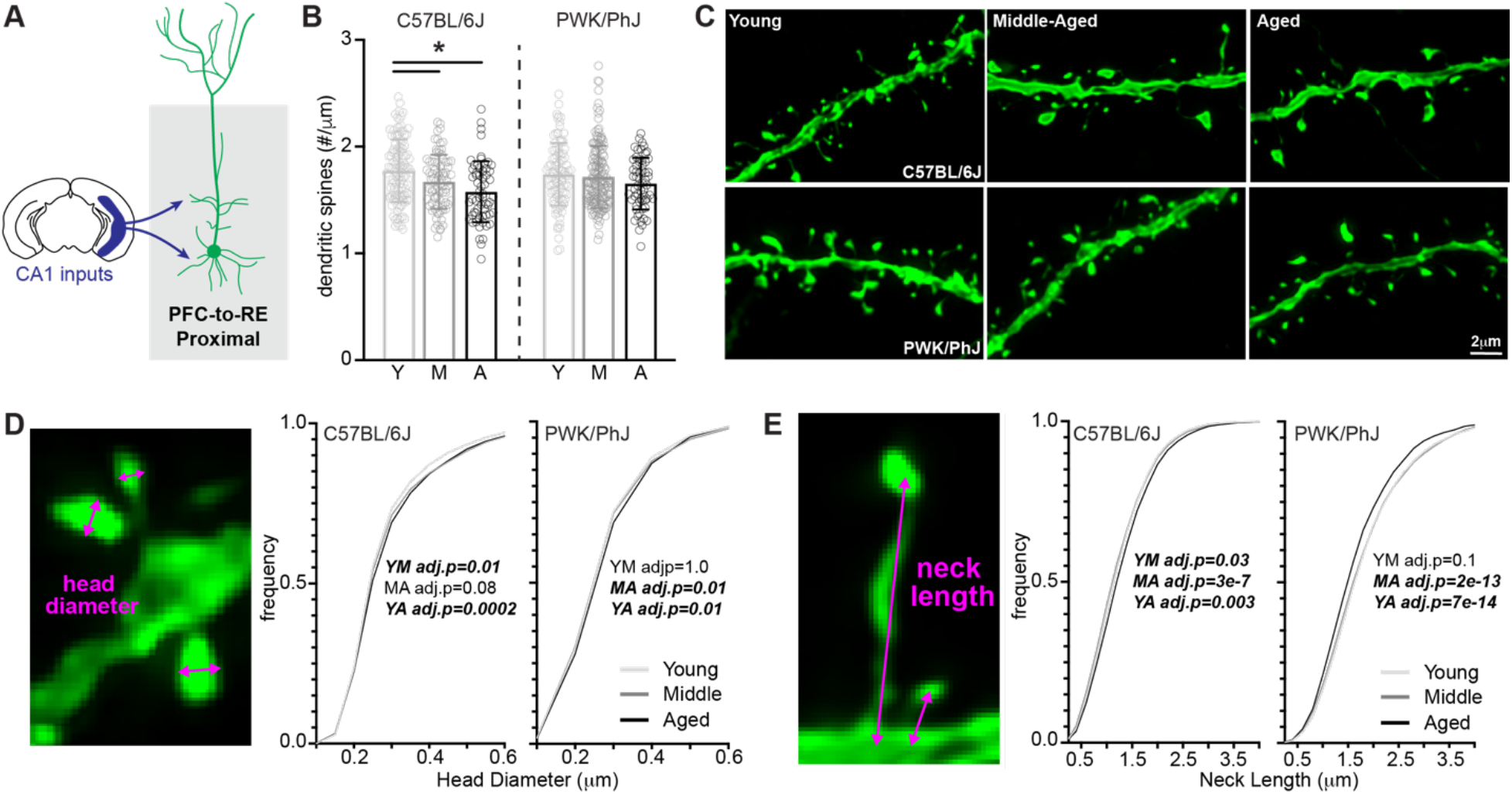
C57BL/6J mice are susceptible to age-related synaptic loss in proximal PFC-to-RE dendrites, with PWK/PhJ demonstrating dynamic morphological changes to increase synaptic strength. **(A)** Example schematic of EGFP+ proximal PFC-to-RE dendrites analyzed here and projection input origins in hippocampal area CA1. **(B)** Proximal PFC-to-RE dendrite spine densities (spines/μm) comparing young (Y), middle-aged (M) and aged (A) within each strain. Data points represent individual branches (n=20/mouse); error bars are ± SD; one-way ANOVA identified significant effect within B6 but not PWK; asterisks denote significant (Bonferroni adjusted p<0.05) comparisons identified using post-hoc analysis within B6 (**Table S4**). **(C)** Representative 63X images of proximal PFC-to-RE dendrites from each strain/age group. **(D)** PFC-to-RE proximal spine head diameter example image (left) and cumulative distributions from B6 (middle) and PWK (right), across age groups. Kolmogorov-Smirnov tests were used to evaluate statistical significance (Bonferroni adj. p<0.05) of pairwise comparisons: young vs middle (YM), middle vs aged (MA), and young vs. aged (YA), and p-values reported on each graph (see **Table S4**). Data points are representative of measures from individual spines. **(E)** Same as **(D)** for PFC-to-RE proximal spine neck length. Summary statistics for data points represented in each graph reported in **Table S4.**

We first examined spine density on proximal dendrites of PFC-to-RE neurons. Consistent with previous findings, B6 proximal dendritic spine density was significantly lower in middle-aged and aged compared to young mice (66). Interestingly, we found no differences in spine densities on proximal PFC dendrites across age cohorts in PWK mice (**Figure 4B-C**). When we examined spine morphologies, we found statistically significant changes in spine head diameter with age on both B6 and PWK proximal PFC dendrites, evidenced by the cumulative frequency analyses and by quartile analysis of the largest and smallest spines (**Figure 4D**, **Figure S3B**). In both B6 and PWK mice, PFC proximal spine head diameters became subtly larger with aging (**Figure 4D**). Interestingly, spines on proximal dendrites from B6 mice showed minimal age-related changes in spine neck length. However, spines on proximal branches from PWK mice were significantly shortened with age, changes that should reduce neck-related filtering of synaptic currents, mimicking the same patterns found on aging CA1 dendrites from PWK mice (**Figure 4E**, **Figure S3C**).

### Age-related dendritic spine loss and morphological remodeling on distal PFC dendrites differs between B6 and PWK mice

Distal tuft branches from PFC-to-RE projection neurons receive distinct afferent inputs compared to proximal branches, coming from the thalamus instead of CA1 (43) (**Figure 5A**). Similar to the age-related changes found on proximal dendrites of PFC-to-RE neurons, distal dendritic spine densities were lower in middle-aged and aged compared to young B6 mice but did not change across age groups in PWK mice (**Figure 5B-C**).

**Figure 5:**
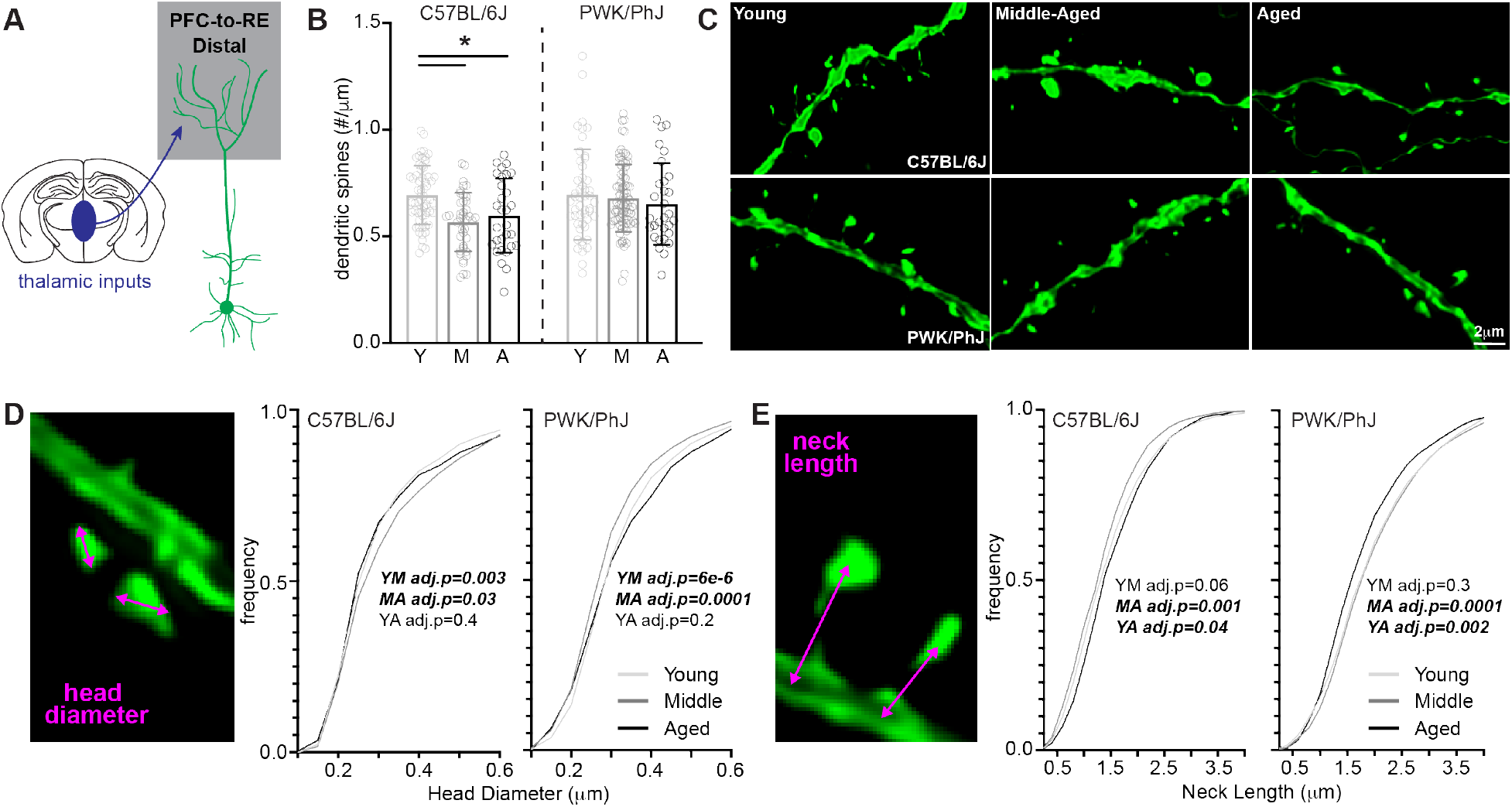
PFC-to-RE distal tuft dendrites exhibit age-induced synaptic loss in C57BL/6J mice but are morphology dynamic in PWK/PhJ mice. **(A)** Example schematic of EGFP+ distal tuft PFC-to-RE dendrites analyzed here and thalamic projection input origins. **(B)** Distal tuft PFC-to-RE dendrite spine densities (spines/μm) comparing young (Y), middle (M) and aged (A) within each strain. Data points represent individual branches (n=10/mouse); error bars are ± SD; one-way ANOVA identified significant effect within B6 but not PWK; asterisks denote significant (Bonferroni adjusted p<0.05) comparisons identified using post-hoc analysis within B6 (**Table S5**). **(C)** Representative 63X images of distal tuft PFC-to-RE dendrites from each strain/age group. **(D)** PFC-to-RE distal tuft spine head diameter example image (left) and cumulative distributions from B6 (middle) and PWK (right), across age groups. Kolmogorov-Smirnov tests were used to evaluate statistical significance (Bonferroni adj. p<0.05) of pairwise comparisons: young vs middle (YM), middle vs aged (MA), and young vs. aged (YA), and p-values reported on each graph (see **Table S5**). Data points are representative of measures from individual spines. **(E)** Same as **(D)** for PFC-to-RE distal tuft spine neck length. Summary statistics for data points represented in each graph reported in **Table S5.**

Spine head diameters on distal branches of PFC-to-RE neurons from B6 mice showed a biphasic response, first increasing in size between young and middle-aged cohorts, then decreasing in size between middle-aged to aged groups. PWK distal PFC spine heads displayed the opposite pattern: decreasing in size between young and middle-aged groups, and then increasing in size between middle-aged and aged cohorts (**Figure 5D, Figure S4B**). Spine neck lengths on the distal dendrites of PFC-to-RE neurons from B6 mice slightly lengthened with age, while those from PWK decreased more dramatically with age (**Figure 5E**). These changes in spine neck lengths were observed in quartile analyses, with a significant gain of long spine necks in aged B6 mice, and an overt loss of long spine necks in aged PWK mice (**Figure S4C**). These results from distal PFC-to-RE dendrites mirrored those from proximal dendrites: B6 branches showed age-related synaptic loss while those from PWK did not. In contrast, PWK distal PFC spines showed spine neck remodeling while those from B6 were markedly more stable.

Collectively, these data support our hypothesis that, like resilience to AD-related Aβ pathology (38), PWK mice show resistance to age-related cortical synapse loss. This resistance was evident across both proximal and distal compartments and accompanied by specific forms of spine head and neck remodeling. Decreases in spine neck length should serve to lower the spine neck resistor and effectively increase the efficiency by which current generated at the synapse reaches the parent dendrite. Given the growing links between dendritic mechanisms of integration and the plasticity that mediates learning and memory (67–70), the putative adaptive spine responses could serve as mechanisms to promote successful maintenance of neuronal health and cognition during aging.

## DISCUSSION

Here, we asked whether the effects of aging on synapse number and structure in two specific neural circuits was equivalent across two distinct strains of mice. We chose to investigate B6 and PWK strains as our past work has identified them as susceptible or resilient to AD-related cognitive and synaptic deficits, respectively (38, 40, 41). To rigorously compare across defined neural circuits, we used multisite viral injections to label CA1-to-PFC and PFC-to-RE neurons and systematically sampled ∼90,000 spines in proximal and distal dendritic compartments across multiple ages (ranging from 5-30 months of age). We found that PFC-projecting CA1 pyramidal cells from both strains maintain their dendritic spine synaptic densities with age, consistent with a body of work examining randomly labeled neurons or synapses in aging NHPs, rats and B6 mice (16, 18, 27, 28, 45). Conversely, we observed a differential pattern of synapse loss on RE-projecting PFC neurons between B6 and PWK mice, with synapse loss restricted to B6 but not PWK mice. Spine reconstructions suggested that spines from B6 showed either stability or patterns of remodeling putatively associated with decreased synaptic efficacy, whereas PWK spines tended to show morphological changes consistent with increased synaptic efficacy during aging (**Table S6**). These results suggest that previously observed cognitive characteristics from these strains (38), including PWK resistance to aging-related decline in cognition or resilience to AD-related neuropathology (38) and differential microglia activation (71) may be due in part to an adaptive forms of synaptic plasticity that maintain connectivity and excitability in cortical circuitry. The PWK mouse strain, then, represents an opportunity to identify and harness adaptive mechanisms to lengthen cognitive resistance to the normal process of aging.

The high-resolution spine reconstructions performed here and viral-based strategies allowed for specific populations of projection neurons to be compared across mice (42). Past work has relied on disparate techniques, including strategies to label random neurons with dye-filling methods or counting random synapses by EM, to build a coherent picture of synaptic vulnerability of the aging brain (22, 25, 29, 35). By taking advantage of multisite viral strategies to gain projection neuron specificity within the same aging subjects, our results are the beginning of identifying the neural circuits that are vulnerable to age-related disconnection in terms of information flow. Interplay between CA1 of the hippocampus, PFC and RE is thought to be critical for spatial working memory tasks, a cognitive domain affected in aged individuals (44, 65, 72). Since our most striking finding is the strain-dependent loss on aging PFC-to-RE neurons, our results suggest the PFC may be the critical locus of aging in terms of synapse loss. Reductions in PFC-to-RE spine densities in B6 mice was most dramatic from young to middle-aged, which is consistent with previous findings of synapses on PFC neurons from rats (25, 30), as well as with behavioral findings that animals and humans start to become impaired in working memory function tasks as early as middle-age (31, 65, 73, 74). Although CA1 projection neurons appear to maintain their numbers during aging, our results do not imply that CA1 function is intact during aging. Other processes, including activity-dependent spine plasticity at CA1 synapses, might also contribute to age-related deficits in hippocampal-dependent learning or memory function.

At the level of individual synapses, spine structure is a strong determinant of function (8, 12, 13). Specifically, spine head diameter is a correlate of synapse strength, and spine neck of the level of compartmentalization between the synapse and parent dendrite. Past work has focused on age-related modification of head diameter with little evidence for modulation of spine neck morphology (62). In contrast, our data suggest that both spine head diameter and neck length are readily modified during aging in a way that depends on genetic context. In B6 mice, CA1-to-PFC neurons exhibited very few morphological spine changes with age, consistent with a remarkable population stability during aging. Conversely, PWK mice showed consistent evidence for morphological spine remodeling on CA1-to-PFC and PFC-to-RE neurons that are suggestive of an overall increase in efficacy between the spine synapse and parent dendrite (**Table S6**). These data are compelling because both spine head and spine neck changes move in unison towards greater synapse efficiency, and their synergism might potentially drive large changes in how excitatory postsynaptic potentials from multiple synapses are integrated in the dendrites.

Recent work from our group has found that PWK mice are resilient to AD-related synaptic changes in CA1-to-PFC neurons during early exponential phases of pathology spread (40). As biological age is the primary risk factor for developing neurodegenerative disorders including AD (75, 76), the data presented here suggest that the PWK mouse strain can be both a model of resilience to cognitive and synaptic changes seen in AD (38, 40), and resistance to cortical synapse loss seen during normal brain aging. Since the molecular and cellular mechanisms that differentiate successful from unsuccessful agers are still unknown (27), the findings here could drive the identification of cell-specific processes that maintain synapses throughout aging. Future work should seek to advance this goal by testing additional PFC projection neurons to understand the circuit specificity of resistance, by molecular strategies to uncover the gene(s) required for this resistance, and by using functional studies that interrogate how behavior relates to pattens of neuronal activity in these circuits (64, 77). Ultimately, such work would set the stage for developing precise therapeutic strategies to promote healthy cognitive aging and resilience to AD across our genetically diverse and aging human population.

## MATERIALS AND METHODS

### Ethics Statement

All research was approved by the Institutional Animal Care and Use Committee (IACUC) at The Jackson Laboratory (approval number 12005 and 20006). Animals were humanely euthanized with 4% tribromoethanol (800 mg/kg). Authors performed their work following guidelines established by “The Eight Edition of the Guide for the Care and Use of Laboratory Animals” and euthanasia using methods approved by the American Veterinary Medical Association.

### Animal husbandry

All mice were bred and housed in a 12/12 hour light/dark cycle on aspen bedding and fed a standard 6% Purina 5K52 Chow diet. Experiments were performed using two mouse strains: C57BL/6J (B6, JAX stock #000664) and PWK/PhJ (PWK, JAX stock #003715). Mice were group housed for entirety of experiments. Experimental cohorts were generated through intercrossing B6 or PWK mice to produce 3-8 female mice per age group. Full mouse information is reported in **Table S1**.

### Intracranial viral injections

Recombinant adenoassociated viral (AAV) vectors were used to drive Cre-recombinase (AAVretro-Cre)(42), Cre-dependent EGFP (serotype 2/1, AAV-FLEX-*rev*-EGFP) (42, 43), Flp-recombinase (AAVretro-CAG-FLPo), and FLP-dependent tdTomato (serotype 2/1 AAV-CAG-FRT-*rev*-TdTomato) (78). AAVretro-CAG-FLPo and AAV-CAG-FRT-*rev*-TdTomato were gifts from Janelia Viral Tools (Addgene plasmid #183412, http://n2t.net/addgene:183412, RRID:Addgene_183412; Addgene plasmid #191203, http://n2t.net/addgene:191203, RRID:Addgene_191203). The titers of each virus were as follows (in genomic copies/mL): AAVretro-Cre, 1×10^12^, AAV-FLEX-*rev*-GFP, 1×10^13^, AAVretro-CAG-FLPo, 1×10^13^; AAV-CAG-FRT-*rev*-TdTomato, 1×10^13^. A 1:1 ratio of AAVretro-CAG-FLPo and AAV-FLEX-*rev*-GFP was mixed and 45nL injected into ventral prefrontal cortex (PFC) over 5 minutes; 45-50 nL (per each D/V coordinate) of AAV-CAG-FRT-*rev*-TdTomato was injected in CA1 (CA1) over 10 minutes; AAVretro-Cre was diluted 1:10 with sterile saline and 45 nL injected into nucleus reunions (RE) over 5 minutes. Since PWK brain volumes are smaller than B6, injection coordinates were adjusted based on pilot experiments to determine injection sites. The coordinates for each injection were as follows (in mm: posterior relative to bregma, lateral relative to midline, and ventral relative to pial surface): B6 PFC (+1.75, -0.95, and -2.6), B6 CA1 (-3.5, -3.4, and -2.7/- 2.5/2.0), B6 RE(-1.1, -1.2, -4.15); PWK PFC (+1.45, -0.9, and -2.3), PWK CA1 (-3.5, -3.3, and -2.75/-2.5/-2.0), PWK RE(-1.0, -1.0, -3.2). RE and PFC injections were performed with the mouse tilted at a 15° angle. At each site the injection pipette was left in place for 3-5 minutes then slowly retracted at a rate of 10 μm/s from the brain. After surgery mice were singly housed, monitored for 5 days, and euthanized approximately 3-4 weeks post injection.

### Tissue harvest and brain sectioning

Mice were euthanized with an intraperitoneal lethal dose of Tribromoethanol (800 mg/kg), followed by transcardial perfusion with 45 mL ice-cold 4% paraformaldehyde (PFA) in 0.1M phosphate-buffered saline, in accordance with IACUC protocols (12005 and 20008). Brains were removed and placed in 5mL ice cold 4% PFA at 4°C for 24 hours, then placed into storage buffer (1XPBS + 0.1% Sodium Azide) for long-term storage at 4°C. Brains were coronally sectioned at 50μm thickness and placed in storage buffer at 4°C until needed for imaging. Approximately 6 frontal PFC sections containing EGFP+ dendrites, and 6 sections with distal CA1 TdTomato+ dendrites were mounted on slides and coverslipped using Vectashield Hardset mounting media (Vector Laboratories #H140010).

### Dendritic spine imaging and analysis

Slices were imaged on an SP8 confocal microscope equipped with a 63X objective (oil immersion), images collected at 50 nm pixel sizes with 0.1 μm z-steps, and stacks deconvolved using Leica Lightning software. 10 dendrites per compartment (e.g. basal, apical oblique, tuft) were captured per mouse. Each image was exported as TIFF format (in ImageJ version 2.9.0/1.53t) and imported into NeuronStudio (79) for analysis of dendritic spine densities and morphologies. Since the effectiveness of a synaptic input on the dendritic tree is influenced by the diameter of the parent dendrite (80), we ensured all dendritic calibers incorporated into this analysis were thin (i.e. under 1 μm in diameter) (**Figure S1A, S2A, S3A, S4A**). Density measurements were acquired by first reconstructing the dendritic cable followed by semi-automated spine identification. Cumulative distributions of assigned spine head diameters (HEAD.DIAMETER) and neck lengths (MAX.DTS) were analyzed by Kolmogorov-Smirnov tests, and through a quartile-based analysis. In this latter analysis, spines within dendritic compartments from each strain were pooled across treatment groups to create a population, and the first and last quartiles determined. From each branch, spines belonging to the first quartile (Q1, smallest) and last quartile (Q4, largest) were identified and the density for each quartile per branch was calculated. Data were analyzed with each dendrite representing an individual data point.

### Statistical analysis

Data were analyzed blinded to strain and age group. All statistical analyses were performed in GraphPad Prism software (v9.5.1) except for Kolmogorov-Smirnov tests which were performed using R (v4.2.2). Results are reported in table form in the Supplemental Information **Table S2- S5**). Data from B6 and PWK mouse strains were analyzed separately. To assess age effects within each strain, one-way ANOVAs were computed followed by Bonferroni post hoc tests. Within-group differences between Q1 and Q4 densities for spine head diameter and neck length were determined using nonparametric two-tailed t-tests. Bonferroni corrections for multiple comparisons were performed on nominal p-values from resultant K-S tests.

### Data and resource sharing plan

All mouse strains are available through The Jackson Laboratory. All reagents in this study are commercially available. Raw data (DOI: 10.6084/m9.figshare.23620752) and images from the figures (DOI: 10.6084/m9.figshare.23620755) are available via Figshare (made public upon acceptance of publication).

### Author contributions

S.E.H., E.B.B. and G.R.H. designed the study. S.E.H. generated and maintained mouse cohorts, and S.E.H. and E.B.B. performed the intracranial AAV injections. S.E.H. euthanized mice and collected tissues. S.E.H. and E.W.N. sectioned brains and prepared slides for imaging. S.E.H. and E.W.N. imaged slices and analyzed final data. E.B.B. provided training in intracranial injections, confocal microscopy imaging of dendrites and dendritic spines, and NeuronStudio analysis. E.B.B. and G.R.H. advised on all data analysis, data interpretation and manuscript preparation. S.E.H., E.W.N., E.B.B. and G.R.H. wrote and edited the manuscript. All authors approved the final version.

## Acknowledgements

We thank Dr. Kristen Onos and Kelly Keezer for providing aged PWK/PhJ mouse cohorts, and Amanda Hewes and Melanie-Maddox Goodrich for providing critical laboratory support for the initiation of these experiments. We also thank Dr. Philipp Henrich at the Microscopy core at The Jackson Laboratory for training and assistance on the confocal microscope.

