## Supplemental Information for "Genetic context drives age-related disparities in synaptic maintenance and structure across cortical and hippocampal neuronal circuits"

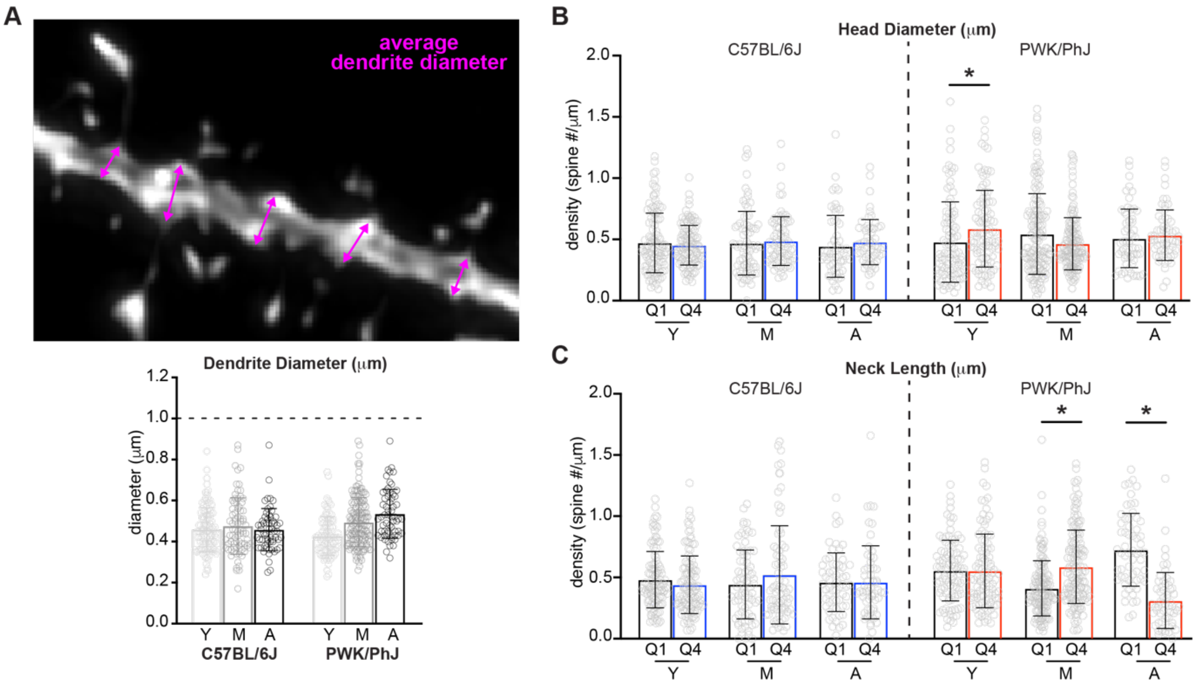


**Figure S1: Additional morphological analyses for proximal CA1-to-PFC spines and dendrites, corresponding to Figure 2.**

**(A)** CA1-to-PFC proximal average dendrite diameter example image (top) and quantification of average diameter for each analyzed dendrite (bottom). Horizontal dashed line represents that dendrites sampled were under 1μm in diameter, corresponding to uniformly thin branches. Data points represent individual branches (n=20/mouse). Groups measured include young (Y), middle-aged (M) and aged (A).

**(B)** Quartile-based analyses of CA1-to-PFC proximal spine head diameters. All proximal CA1-to-PFC spines within each strain were divided into quartiles based on head diameter (μm). The smallest spines assigned to the first quartile (Q1) and the largest spines assigned to the fourth quartile (Q4) were identified and reassigned back to originating dendrite. Spine densities (spines/μm) for Q1 and Q4 spines were calculated separately. Data points represent individual branches. Nonparametric two-tailed t-tests were performed to compare Q1 to Q4 within each strain/age group to identify significant (*=p<0.05) shifts in size (see **Table S2**).

**(C)** Same as **(B)** for CA1-to-PFC proximal spine neck length.

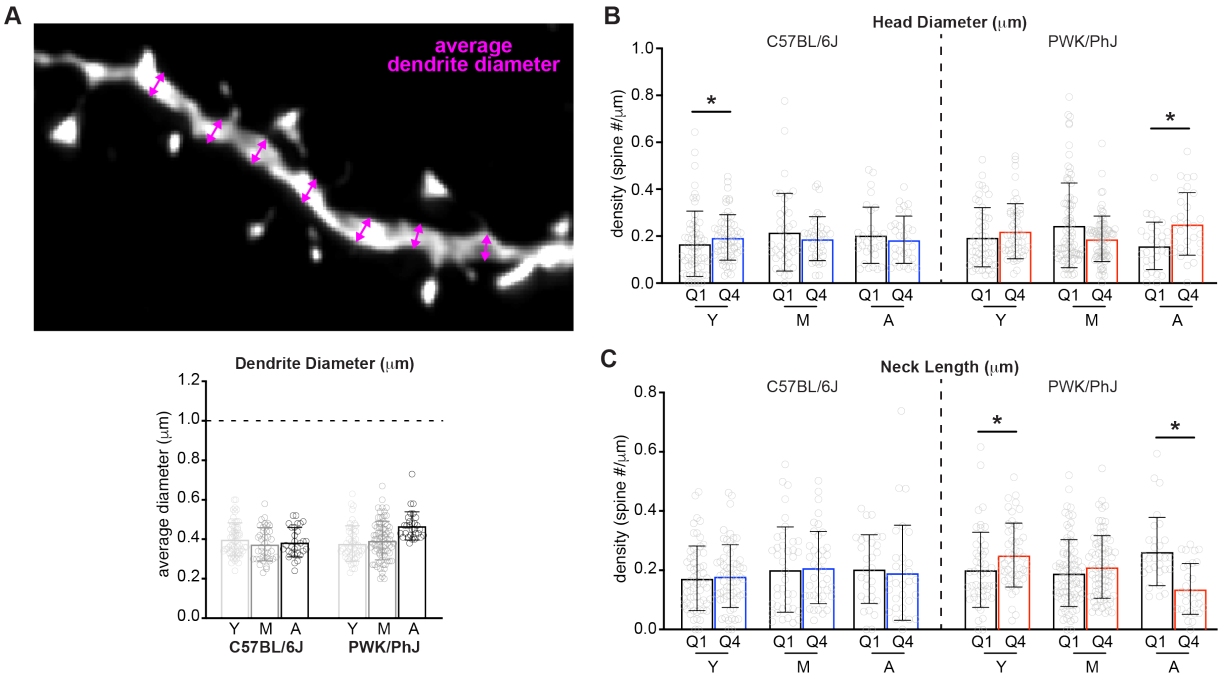


**Figure S2: Additional morphological analyses for distal tuft CA1-to-PFC spines and dendrites, corresponding to Figure 3.**

**(A)** CA1-to-PFC distal average dendrite diameter example image (top) and quantification of average diameter for each analyzed dendrite (bottom). Horizontal dashed line represents that dendrites sampled were under 1μm in diameter, corresponding to uniformly thin branches. Data points represent individual branches (n=10/mouse). Groups measured include young (Y), middle-aged (M) and aged (A).

**(B)** Quartile-based analyses of distal tuft CA1-to-PFC spine head diameters. All distal tuft CA1-to-PFC spines within each strain were divided into quartiles based on head diameter (μm). The smallest spines assigned to the first quartile (Q1) and the largest spines assigned to the fourth quartile (Q4) were identified and reassigned back to originating dendrite. Spine densities (spines/μm) for Q1 and Q4 spines were calculated separately. Data points represent individual branches. Nonparametric two-tailed t-tests were performed to compare Q1 to Q4 within each strain/age group to identify significant (*=$p<0.05$) shifts in size (see **Table S3**).

**(C)** Same as **(B)** for CA1-to-PFC distal tuft spine neck length.

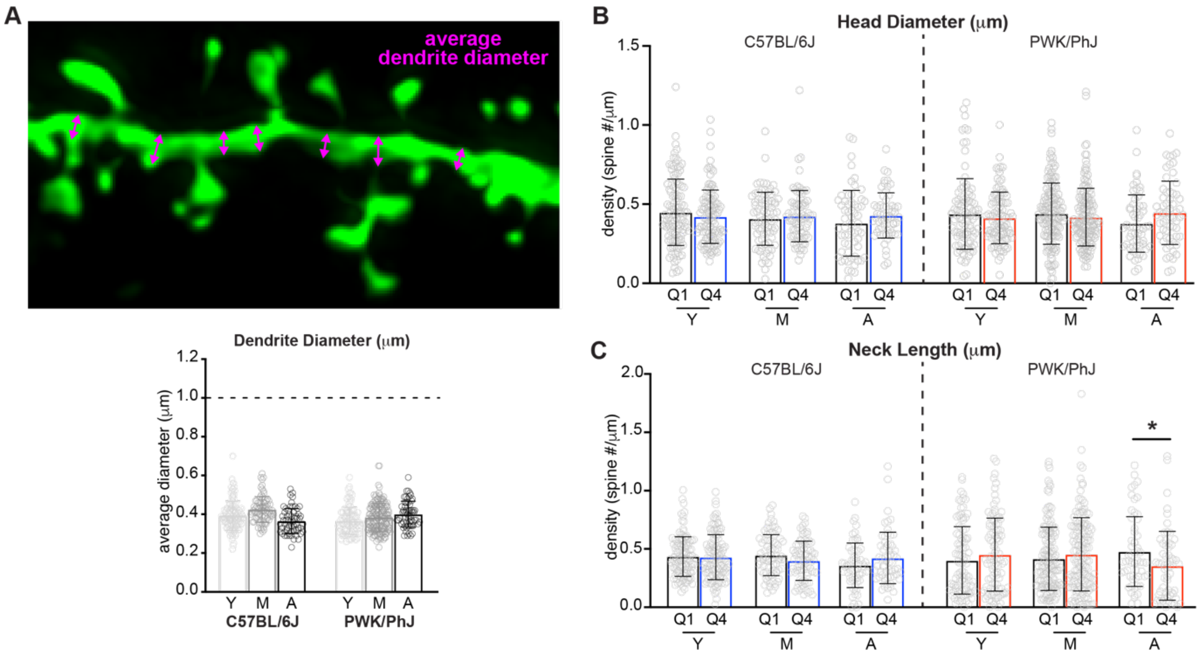


**Figure S3: Additional morphological analyses for distal tuft PFC-to-RE spines and dendrites, corresponding to Figure 4.**

**(A)** PFC-to-RE proximal average dendrite diameter example image (top) and quantification of average diameter for each analyzed dendrite (bottom). Horizontal dashed line represents that dendrites sampled were under 1μm in diameter, corresponding to uniformly thin branches. Data points represent individual branches (n=20/mouse). Groups measured include young (Y), middle-aged (M) and aged (A).

**(B)** Quartile-based analyses of proximal PFC-to-RE spine head diameters. All proximal PFC-to-RE spines within each strain were divided into quartiles based on head diameter (μm). The smallest spines assigned to the first quartile (Q1) and the largest spines assigned to the fourth quartile (Q4) were identified and reassigned back to originating dendrite. Spine densities (spines/μm) for Q1 and Q4 spines were calculated separately. Data points represent individual branches. Nonparametric two-tailed t-tests were performed to compare Q1 to Q4 within each strain/age group to identify significant (*=p<0.05) shifts in size (see **Table S4**).

**(C)** Same as **(B)** for PFC-to-RE proximal spine neck length.

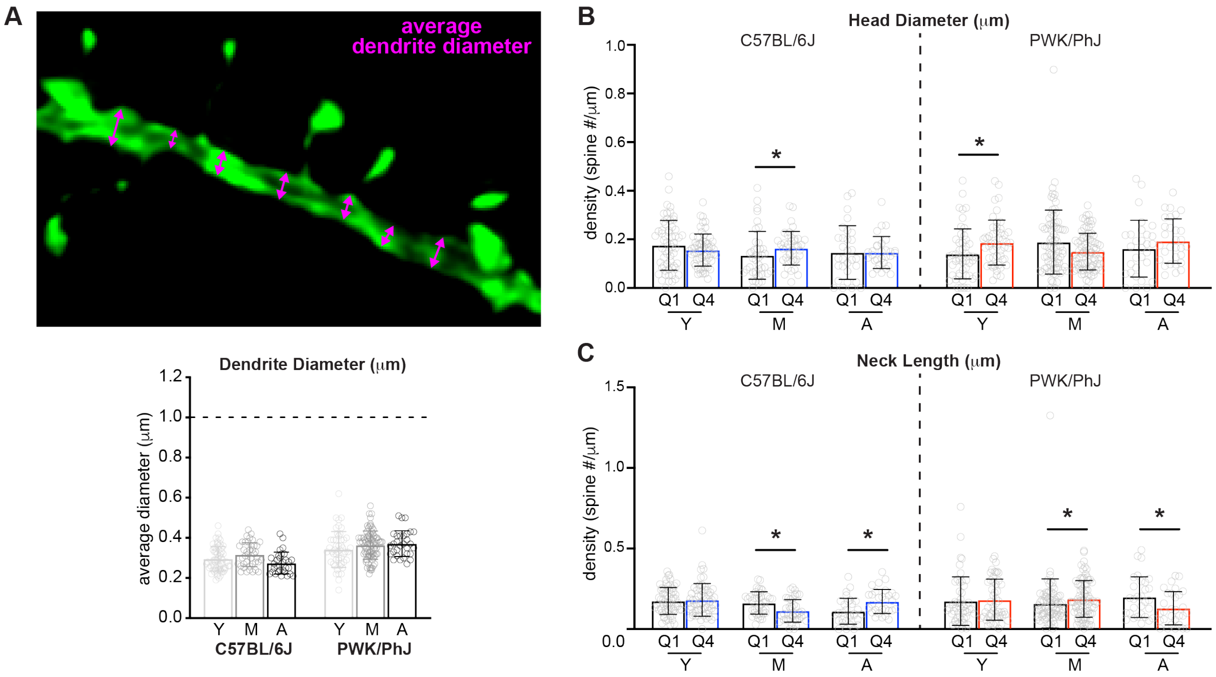


**Figure S4: Additional morphological analyses for distal tuft PFC-to-RE spines and dendrites, corresponding to Figure 5.**

**(A)** PFC-to-RE distal average dendrite diameter example image (top) and quantification of average diameter for each analyzed dendrite (bottom). Horizontal dashed line represents that dendrites sampled were under 1μm in diameter, corresponding to uniformly thin branches. Data points represent individual branches (n=10/mouse). Groups measured include young (Y), middle-aged (M) and aged (A).

**(B)** Quartile-based analyses of distal tuft PFC-to-RE spine head diameters. All distal tuft PFC-to-RE spines within each strain were divided into quartiles based on head diameter (μm). The smallest spines assigned to the first quartile (Q1) and the largest spines assigned to the fourth quartile (Q4) were identified and reassigned back to originating dendrite. Spine densities (spines/μm) for Q1 and Q4 spines were calculated separately. Data points represent individual branches. Nonparametric two-tailed t-tests were performed to compare Q1 to Q4 within each strain/age group to identify significant (*=$p<0.05$) shifts in size (see **Table S5**).

**(C)** Same as **(B)** for PFC-to-RE distal tuft spine neck length.

### SUPPLEMENTAL TABLES

#### Table S1- Mouse information for current study, corresponding to Figure 1

| MouseID | Strain | Sex | DOB | Harvest Date | Age (m) | Age Group | Terminal body weight (g) |
|---|---|---|---|---|---|---|---|
| 59944 | PWK/PhJ | F | 11/24/2019 | 5/21/2022 | 30 | Aged | 18.4 |
| 59246 | PWK/PhJ | F | 4/24/2020 | 5/21/2022 | 25 | Aged | 24.6 |
| 59247 | PWK/PhJ | F | 4/24/2020 | 5/21/2022 | 25 | Aged | 23.3 |
| 76266 | PWK/PhJ | F | 3/9/2021 | 8/8/2022 | 17 | Middle-Aged | 17.1 |
| 76267 | PWK/PhJ | F | 3/9/2021 | 8/8/2022 | 17 | Middle-Aged | 21.6 |
| 60938 | PWK/PhJ | F | 3/16/2021 | 6/13/2022 | 15 | Middle-Aged | 19.4 |
| 60939 | PWK/PhJ | F | 3/16/2021 | 6/13/2022 | 15 | Middle-Aged | 23.9 |
| 60942 | PWK/PhJ | F | 3/16/2021 | 6/13/2022 | 15 | Middle-Aged | 18.1 |
| 56621 | PWK/PhJ | F | 6/23/2021 | 6/30/2022 | 12 | Middle-Aged | 21.1 |
| 56622 | PWK/PhJ | F | 6/23/2021 | 6/30/2022 | 12 | Middle-Aged | 18.6 |
| 56833 | PWK/PhJ | F | 7/14/2021 | 6/30/2022 | 11 | Middle-Aged | 17.2 |
| 70328 | PWK/PhJ | F | 11/5/2021 | 6/13/2022 | 7 | Young | 16.3 |
| 70329 | PWK/PhJ | F | 11/5/2021 | 6/13/2022 | 7 | Young | 18.0 |
| 27811R | PWK/PhJ | F | 1/11/2022 | 6/13/2022 | 5 | Young | 16.1 |
| 2781B | PWK/PhJ | F | 1/11/2022 | 6/13/2022 | 5 | Young | 17.6 |
| 27812L | PWK/PhJ | F | 1/11/2022 | 6/13/2022 | 5 | Young | 16.9 |
| 2430A | C57BL/6J | F | 8/9/2022 | 12/13/2022 | 4 | Young | 21.1 |
| 2430B | C57BL/6J | F | 8/9/2022 | 12/13/2022 | 4 | Young | 21.0 |
| A0003 | C57BL/6J | F | 5/31/2022 | 11/4/2022 | 6 | Young | 25.7 |
| A0004 | C57BL/6J | F | 5/31/2022 | 11/4/2022 | 6 | Young | 23.8 |
| 24311R | C57BL/6J | F | 5/10/2022 | 12/18/2022 | 7 | Young | 23.9 |
| 2432A | C57BL/6J | F | 5/10/2022 | 12/18/2022 | 7 | Young | 27.6 |
| 5551R | C57BL/6J | F | 11/23/2021 | 12/18/2022 | 13 | Middle-Aged | 25.3 |
| 555B | C57BL/6J | F | 11/23/2021 | 12/13/2022 | 13 | Middle-Aged | 26.1 |
| 15721R | C57BL/6J | F | 7/13/2021 | 12/13/2022 | 17 | Middle-Aged | 27.4 |
| 15722R | C57BL/6J | F | 7/13/2021 | 12/18/2022 | 17 | Middle-Aged | 36.9 |
| 60840 | C57BL/6J | F | 2/16/2021 | 12/13/2022 | 22 | Aged | 35.7 |
| A3924 | C57BL/6J | F | 12/2/2020 | 11/4/2022 | 23 | Aged | 37.9 |
| A3926 | C57BL/6J | F | 12/2/2020 | 12/18/2022 | 24 | Aged | 29.1 |

### Table S2- Associated statistics for proximal CA1 dendrites, corresponding to Figure 2 and Figure S1

#### S2A. Summary statistics for proximal CA1-to-PFC spine data

| Strain | Age Group | Mouse N | Dendrite N | Spine N |
|---|---|---|---|---|
| B6 | Young | 6 | 120 | 7479 |
| B6 | Middle-Aged | 4 | 80 | 4845 |
| B6 | Aged | 3 | 60 | 3611 |
| PWK | Young | 5 | 100 | 7125 |
| PWK | Middle-Aged | 8 | 160 | 10882 |
| PWK | Aged | 3 | 60 | 4494 |

#### S2B. Spine density ANOVA followed by Bonferroni post-hoc pairwise analysis

| Group A | Group B | C57BL/6J Adj. p-value | PWK/PhJ Adj. p-value |
|---|---|---|---|
| Young | Middle-Aged | >0.9999 | >0.9999 |
| Middle-Aged | Aged | 0.9088 | 0.6156 |
| Young | Aged | 0.2297 | >0.9999 |
| One-way ANOVA | | F = 1.584<br>p-value = 0.2071 | F = 0.8171<br>p-value = 0.4427 |

#### S2C. Head Diameter Kolmogorov-Smirnov tests

| | | C57BL/6J | | PWK/PhJ | |
|---|---|---|---|---|---|
| Group A | Group B | KS p-value | K-S Bonferroni adjusted pvalue | KS p-value | K-S Bonferroni adjusted pvalue |
| Young | Middle-Aged | 0.02206 | 0.06618 | 2.2e-16 | 6.6e-16 |
| Middle-Aged | Aged | 0.07032 | 0.21096 | 0.006142 | 0.018426 |
| Young | Aged | 0.01216 | 0.03648 | 3.542e-07 | 1.0626E-06 |

#### S2D. Head diameter quartile nonparametric t-tests

| Group A | Group B | C57BL/6J p-value | PWK/PhJ p-value |
|---|---|---|---|
| Young Q1 | Young Q4 | 0.8133 | 0.0030 |
| Middle-Aged Q1 | Middle-Aged Q4 | 0.2422 | 0.1786 |
| Aged Q1 | Aged Q4 | 0.0715 | 0.3116 |

#### S2E. Neck Length Kolmogorov-Smirnov tests

| | | C57BL/6J | | PWK/PhJ | |
|---|---|---|---|---|---|
| Group A | Group B | KS p-value | K-S Bonferroni adjusted pvalue | KS p-value | K-S Bonferroni adjusted pvalue |
| Young | Middle-Aged | 4.836e-11 | 1.4508E-10 | 2.2e-16 | 6.6e-16 |
| Middle-Aged | Aged | 0.002058 | 0.006174 | 2.2e-16 | 6.6e-16 |
| Young | Aged | 0.02071 | 0.06213 | 2.2e-16 | 6.6e-16 |

#### S2F. Neck length quartile nonparametric t-tests

| Group A | Group B | C57BL/6J p-value | PWK/PhJ p-value |
|---|---|---|---|
| Young Q1 | Young Q4 | 0.1029 | 0.4625 |
| Middle-Aged Q1 | Middle-Aged Q4 | 0.6502 | <0.0001 |
| Aged Q1 | Aged Q4 | 0.5712 | <0.0001 |

### Table S3- Associated statistics for distal tuft CA1 dendrites, corresponding to Figure 3 and Figure S2

#### S3A. Summary statistics for distal CA1-to-PFC spine data

| Strain | Age Group | Mouse N | Dendrite N | Spine N |
|---|---|---|---|---|
| B6 | Young | 6 | 60 | 1370 |
| B6 | Middle-Aged | 4 | 40 | 1003 |
| B6 | Aged | 3 | 30 | 700 |
| PWK | Young | 5 | 50 | 1512 |
| PWK | Middle-Aged | 8 | 80 | 2123 |
| PWK | Aged | 3 | 30 | 849 |

#### S3B. Spine density ANOVA followed by Bonferroni post-hoc pairwise analysis

| Group A | Group B | C57BL/6J Adj. p-value | PWK/PhJ Adj. p-value |
|---|---|---|---|
| Young | Middle-Aged | >0.9999 | >0.9999 |
| Middle-Aged | Aged | >0.9999 | >0.9999 |
| Young | Aged | >0.9999 | >0.9999 |
| One-way ANOVA | | F = 0.4320<br>p-value = 0.6502 | F = 0.2171<br>p-value = 0.8051 |

#### S3C. Head Diameter Kolmogorov-Smirnov tests

| | | C57BL/6J | | PWK/PhJ | |
|---|---|---|---|---|---|
| Group A | Group B | KS p-value | K-S Bonferroni adjusted pvalue | KS p-value | K-S Bonferroni adjusted pvalue |
| Young | Middle-Aged | 0.004019 | 0.012057 | 0.0004517 | 0.0013551 |
| Middle-Aged | Aged | 0.1044 | 0.3132 | 4.668e-09 | 1.4004E-08 |
| Young | Aged | 0.2682 | 0.8046 | 0.005925 | 0.017775 |

#### S3D. Head diameter quartile nonparametric t-tests

| Group A | Group B | C57BL/6J p-value | PWK/PhJ p-value |
|---|---|---|---|
| Young Q1 | Young Q4 | 0.0382 | 0.1527 |
| Middle-Aged Q1 | Middle-Aged Q4 | 0.9268 | 0.2656 |
| Aged Q1 | Aged Q4 | 0.7330 | 0.0060 |

#### S3E. Neck Length Kolmogorov-Smirnov tests

| | | C57BL/6J | | PWK/PhJ | |
|---|---|---|---|---|---|
| Group A | Group B | KS p-value | K-S Bonferroni adjusted pvalue | KS p-value | KS Bonferroni adjusted pvalue |
| Young | Middle-Aged | 0.03513 | 0.10539 | 0.06974 | 0.20922 |
| Middle-Aged | Aged | 0.2409 | 0.7227 | 9.98e-07 | 2.994E-06 |
| Young | Aged | 0.01058 | 0.03174 | 2.762e-08 | 8.286E-08 |

#### S3F. Neck length quartile nonparametric t-tests

| Group A | Group B | C57BL/6J p-value | PWK/PhJ p-value |
|---|---|---|---|
| Young Q1 | Young Q4 | 0.5801 | 0.0089 |
| Middle-Aged Q1 | Middle-Aged Q4 | 0.6170 | 0.1903 |
| Aged Q1 | Aged Q4 | 0.3561 | <0.0001 |

### Table S4- Associated statistics for proximal PFC dendrites, corresponding to Figure 4 and Figure S3

#### S4A. Summary statistics for proximal PFC-to-RE spine data

| Strain | Age Group | Mouse N | Dendrite N | Spine N |
|---|---|---|---|---|
| B6 | Young | 6 | 120 | 7997 |
| B6 | Middle-Aged | 4 | 80 | 5126 |
| B6 | Aged | 3 | 60 | 3419 |
| PWK | Young | 5 | 100 | 6754 |
| PWK | Middle-Aged | 8 | 160 | 10611 |
| PWK | Aged | 3 | 60 | 3853 |

#### S4B. Spine density ANOVA followed by Bonferroni post-hoc pairwise analysis

| Group A | Group B | C57BL/6J<br>Adj. p-value | PWK/PhJ<br>Adj. p-value |
|---|---|---|---|
| Young | Middle-Aged | 0.0339 | >0.9999 |
| Middle-Aged | Aged | 0.1552 | 0.4328 |
| Young | Aged | <0.0001 | 0.1846 |
| One-way ANOVA | | F = 10.36<br>p-value < 0.0001 | F = 1.796<br>p-value = 0.1676 |

#### S4C. Head Diameter Kolmogorov-Smirnov tests

| | | C57BL/6J | | PWK/PhJ | |
|---|---|---|---|---|---|
| Group A | Group B | KS p-value | K-S Bonferroni adjusted pvalue | KS p-value | K-S Bonferroni adjusted pvalue |
| Young | Middle-Aged | 0.003496 | 0.010488 | 0.3364 | 1 |
| Middle-Aged | Aged | 0.02801 | 0.08403 | 0.003807 | 0.011421 |
| Young | Aged | 7.682e-05 | 0.00023046 | 0.00315 | 0.00945 |

#### S4D. Head Diameter quartile nonparametric t-tests

| Group A | Group B | C57BL/6J<br>p-value | PWK/PhJ<br>p-value |
|---|---|---|---|
| Young Q1 | Young Q4 | 0.3153 | 0.7610 |
| Middle-Aged Q1 | Middle-Aged Q4 | 0.8602 | 0.1076 |
| Aged Q1 | Aged Q4 | 0.1300 | 0.0503 |

#### S4E. Neck Length Kolmogorov-Smirnov tests

| | | C57BL/6J | | PWK/PhJ | |
|---|---|---|---|---|---|
| Group A | Group B | KS p-value | K-S Bonferroni adjusted pvalue | KS p-value | K-S Bonferroni adjusted pvalue |
| Young | Middle-Aged | 0.01101 | 0.03303 | 0.03866 | 0.11598 |
| Middle-Aged | Aged | 1.002e-07 | 3.006e-07 | 5.029e-14 | 1.5087e-13 |
| Young | Aged | 0.0009113 | 0.0027339 | 2.343e-14 | 7.029e-14 |

#### S4F. Neck length quartile nonparametric t-tests

| Group A | Group B | C57BL/6J<br>p-value | PWK/PhJ<br>p-value |
|---|---|---|---|
| Young Q1 | Young Q4 | 0.6594 | 0.3307 |
| Middle-Aged Q1 | Middle-Aged Q4 | 0.1251 | 0.3613 |
| Aged Q1 | Aged Q4 | 0.1942 | 0.0179 |

### Table S5- Associated statistics for distal tuft PFC dendrites, corresponding to Figure 5 and Figure S4

#### S5A. Summary statistics for distal PFC-to-RE spine data

| Strain | Age Group | Mouse N | Dendrite N | Spine N |
|---|---|---|---|---|
| B6 | Young | 6 | 60 | 1481 |
| B6 | Middle-Aged | 4 | 40 | 815 |
| B6 | Aged | 3 | 30 | 631 |
| PWK | Young | 5 | 50 | 1321 |
| PWK | Middle-Aged | 8 | 80 | 2085 |
| PWK | Aged | 3 | 30 | 740 |

#### S5B. Spine density ANOVA followed by Bonferroni post-hoc pairwise analysis

| Group A | Group B | C57BL/6J Adj. p-value | PWK/PhJ Adj. p-value |
|---|---|---|---|
| Young | Middle-Aged | 0.0001 | >0.9999 |
| Middle-Aged | Aged | >0.9999 | >0.9999 |
| Young | Aged | 0.0124 | 0.9075 |
| One-way ANOVA | | F = 9.963<br>p-value <0.0001 | F = 0.5352<br>p-value = 0.5866 |

#### S5C. Head Diameter Kolmogorov-Smirnov tests

| | | C57BL/6J | | PWK/PhJ | |
|---|---|---|---|---|---|
| Group A | Group B | KS p-value | K-S Bonferroni adjusted pvalue | KS p-value | K-S Bonferroni adjusted pvalue |
| Young | Middle-Aged | 0.001061 | 0.003183 | 2.009e-06 | 6.027e-06 |
| Middle-Aged | Aged | 0.008744 | 0.026232 | 4.142e-05 | 0.00012426 |
| Young | Aged | 0.1408 | 0.4224 | 0.07482 | 0.22446 |

#### S5D. Head diameter quartile nonparametric t-tests

| Group A | Group B | C57BL/6J p-value | PWK/PhJ p-value |
|---|---|---|---|
| Young Q1 | Young Q4 | 0.3441 | 0.0073 |
| Middle-Aged Q1 | Middle-Aged Q4 | 0.0187 | 0.0944 |
| Aged Q1 | Aged Q4 | 0.8058 | 0.2065 |

#### S5E. Neck Length Kolmogorov-Smirnov tests

| | | C57BL/6J | | PWK/PhJ | |
|---|---|---|---|---|---|
| Group A | Group B | KS p-value | K-S Bonferroni adjusted pvalue | KS p-value | K-S Bonferroni adjusted pvalue |
| Young | Middle-Aged | 0.01932 | 0.05796 | 0.09178 | 0.27534 |
| Middle-Aged | Aged | 0.0003468 | 0.0010404 | 3.574e-05 | 0.00010722 |
| Young | Aged | 0.0131 | 0.0393 | 0.0006769 | 0.0020307 |

#### S5F. Neck length quartile nonparametric t-tests

| Group A | Group B | C57BL/6J p-value | PWK/PhJ p-value |
|---|---|---|---|
| Young Q1 | Young Q4 | 0.9760 | 0.6223 |
| Middle-Aged Q1 | Middle-Aged Q4 | 0.0023 | 0.0375 |
| Aged Q1 | Aged Q4 | 0.0019 | 0.0418 |

### Table S6- Summary of spine responses to age within each strain

#### S6A. CA1 Spine changes: C57BL/6J

| | | Experimental measures | | | Predicted measures | |
|---|---|---|---|---|---|---|
| | Age Groups | Density | Head Diameter | Neck Length | EPSP spine | EPSP dendrite |
| Proximal | Young → Middle-Aged | -- | -- | ↑ | -- | ↓ |
| | Middle-Aged → Aged | -- | -- | ↓ | -- | ↑ |
| | Young → Aged | -- | ↑ | -- | ↑ | -- |
| Distal | Young → Middle-Aged | -- | ↓ | -- | ↓ | -- |
| | Middle-Aged → Aged | -- | -- | -- | -- | -- |
| | Young → Aged | -- | -- | ↓ | -- | ↑ |

#### S6B. CA1 Spine changes: PWK/PhJ

| | | Experimental measures | | | Predicted measures | |
|---|---|---|---|---|---|---|
| | Age Groups | Density | Head Diameter | Neck Length | EPSP spine | EPSP dendrite |
| Proximal | Young → Middle-Aged | -- | ↓ | ↑ | ↓ | ↓ |
| | Middle-Aged → Aged | -- | ↑ | ↓ | ↑ | ↑ |
| | Young → Aged | -- | ↓ | ↓ | ↓ | ↑ |
| Distal | Young → Middle-Aged | -- | ↓ | -- | ↓ | -- |
| | Middle-Aged → Aged | -- | ↑ | ↓ | ↑ | ↑ |
| | Young → Aged | -- | ↑ | ↓ | ↑ | ↑ |

**S6C. PFC Spine changes: C57BL/6J**

| | | Experimental measures | | | Predicted measures | |
|---|---|---|---|---|---|---|
| | Age Groups | Density | Head Diameter | Neck Length | EPSP spine | EPSP dendrite |
| Proximal | Young → Middle-Aged | ↓ | ↑ | ↓ | ↑ | ↑ |
| | Middle-Aged → Aged | -- | -- | ↑ | -- | ↓ |
| | Young → Aged | ↓ | ↑ | ↑ | ↑ | ↓ |
| Distal | Young → Middle-Aged | ↓ | ↑ | -- | ↑ | -- |
| | Middle-Aged → Aged | -- | ↓ | ↑ | ↓ | ↓ |
| | Young → Aged | ↓ | -- | ↑ | -- | ↓ |

**S6D. PFC Spine changes: PWK/PhJ**

| | | Experimental measures | | | Predicted measures | |
|---|---|---|---|---|---|---|
| | Age Groups | Density | Head Diameter | Neck Length | EPSP spine | EPSP dendrite |
| Proximal | Young → Middle-Aged | -- | -- | -- | -- | -- |
| | Middle-Aged → Aged | -- | ↑ | ↓ | ↑ | ↑ |
| | Young → Aged | -- | ↑ | ↓ | ↑ | ↑ |
| Distal | Young → Middle-Aged | -- | ↓ | -- | ↓ | -- |
| | Middle-Aged → Aged | -- | ↑ | ↓ | ↑ | ↑ |
| | Young → Aged | -- | -- | ↓ | -- | ↑ |
